# Physiological and Metabolic Features of Mice with CRISPR/Cas9-Mediated Loss-of-Function in Growth Hormone-Releasing Hormone

**DOI:** 10.1101/2020.02.07.937789

**Authors:** Mert Icyuz, Michael P. Fitch, Fang Zhang, Anil K. Challa, Liou Y. Sun

## Abstract

Our previous study demonstrated that the loss of growth hormone releasing hormone (GHRH) results in increased lifespan and improved metabolic homeostasis in the mouse model generated by classical embryonic stem cell based gene-targeting method. In this study, we targeted the GHRH gene using the CRISPR/Cas9 technology to avoid passenger alleles/mutations and performed in-depth physiological and metabolic characterization. In agreement with our previous observation, male and female GHRH^-/-^ mice have significantly reduced body weight and enhanced insulin sensitivity when compared to wild type littermates. Dual-energy X-ray absorptiometry showed that there were significant decreases in lean mass, bone mineral content and density, and a dramatic increase in fat mass of GHRH^-/-^ mice when compared to wild type littermates. Indirect calorimetry measurements including oxygen consumption, carbon dioxide production and energy expenditure were dramatically reduced in GHRH^-/-^ mice compared to wild type mice. Respiratory exchange ratio was significantly lower in GHRH^-/-^ mice during the light cycle, but not during the dark cycle, indicating a circadian related metabolic shift towards fat utilization in the growth hormone deficient mice. The novel CRISPR/Cas9 GHRH^-/-^ mice are exhibiting the consistent and unique physiological and metabolic characteristics, which might mediate the longevity effects of growth hormone deficiency in mice.

## Introduction

Aging is defined as a progressive decline in physiological function, which results in increased risk of chronic diseases such as cancer, diabetes, and Alzheimer’s [1]. Geroscience has established that aging is influenced by genetic and environmental factors and genetically regulated longevity mechanisms have been identified in *Caenorhabditis elegans, Drosophila melanogaster* and *Mus musculus* [2, 3]. Endocrine signaling is the most studied modulator of longevity in animal models [4–7]. Specifically, suppression of growth hormone (GH) signaling pathway has been linked to increased lifespan in mice [6, 8, 9]. A recent study involving human subjects showed that familial longevity is associated with lower GH secretion [10].

Ames and Snell mice are dwarf due to mutations in the Prophet of Pit-1 (Prop-1) and pituitary factor-1 (Pit1) genes, respectively, in the anterior pituitary [11, 12]. These mutations result in suppression of growth hormone signaling in mice, with delayed aging and improved longevity with increased insulin sensitivity. In addition, they have reduced age-related loss of cognitive function and decreased occurrence of neoplastic disease [13]. In dwarf mice, besides growth hormone deficiency, prolactin (PRL) is absent and levels of thyroid-stimulating hormone (TSH) is greatly reduced in the plasma [14, 15]. GH receptor/GH-binding protein knockout (GHRKO) mice were made in the Kopchick lab to disrupt the Ghr/Ghrbp gene. However, knock out of the Ghr/Ghrbp gene results in dramatic increases of serum growth hormone in both sexes [16].

To generate an isolated growth hormone deficiency model, the Salvatori lab knocked out growth hormone releasing hormone (GHRH), which is a hypothalamic peptide that controls both the synthesis and secretion of GH, in mice using traditional embryonic stem cell (ESC) based gene-targeting method [17, 18]. These mutant mice have significantly decreased body weight and possess increased insulin sensitivity and prolonged lifespan indicating GH deficiency is primarily responsible for longevity extension [5]. This GHRH^-/-^ model was generated using 129SV agouti color mice resulting in the co-segregation of GHRH^-/-^ and agouti alleles. Agouti has shown to be an important component of several biological pathways, including body weight homeostasis, regulation of food intake and, energy expenditure [19]. In addition, its expression was associated with metabolic syndrome [20]. Our goal for this study is to delineate the direct physiological and metabolic consequences of growth hormone deficiency. To achieve this goal, we knocked out the GHRH gene with CRISPR/Cas9 technology, preventing the agouti gene from acting as a passenger allele. In addition, we produced our experimental knockout model on mixed genetic background to avoid any phenotype resulting from strain-specific inbreeding. This is the first in-depth metabolic and physiological profiling of GH-suppressed mice generated with CRISPR/Cas9 technology on mixed genetic background.

Our novel GHRH^-/-^ mice have decreased body weight and higher insulin sensitivity despite having normal glucose tolerance. GH deficiency resulted in dramatically decreased BMD, BMC, and lean mass. However, GHRH^-/-^ mice have significantly increased fat mass compared to littermate controls. Indirect calorimetry allowed us to measure physiological respiratory parameters, which were used to calculate respiratory exchange ratio (RER) and energy expenditure. RER data demonstrated a significant difference in metabolism between GHRH^-/-^ and wild type (WT) mice during light cycle. GHRH^-/-^ mice had significantly lower energy expenditure during both light and dark cycles. Our GH-deficient model has significantly higher insulin sensitivity compared to WT mice. GHRH^-/-^ mice have physiological characteristics consistent with our previous study and similar to other GH related mutants. It was hypothesized slowing the biological process of aging is associated with GH deficiency. We believe our model will help understand key physiological and metabolic characteristics that are involved in the process of aging.

## Results

Utilizing CRISPR/Cas9-mediated gene-editing method, we generated homozygous GHRH^-/-^ mice (Fig. 1A). In a litter of 10 G_0_ pups, 8 carried indels and large deletions in the GHRH locus. One allele (#28528), a 291 base pairs deletion that eliminates the splice donor site at exon 2, intron 2-3 and a large part of Exon 3 (77 base pairs out of 102 base pairs), showed successful germline transmission (Fig. 1A, B). Predicted translation of the resulting sequence suggests an in-frame mutation leading to a loss of 26 amino acids, including 21 amino acids of the minimum sequence required for full activity (RMQRHVDAIFTTNYRKLLSQLYARKV). The line carrying this mutant allele was used in all subsequent studies.

**Figure 1.**
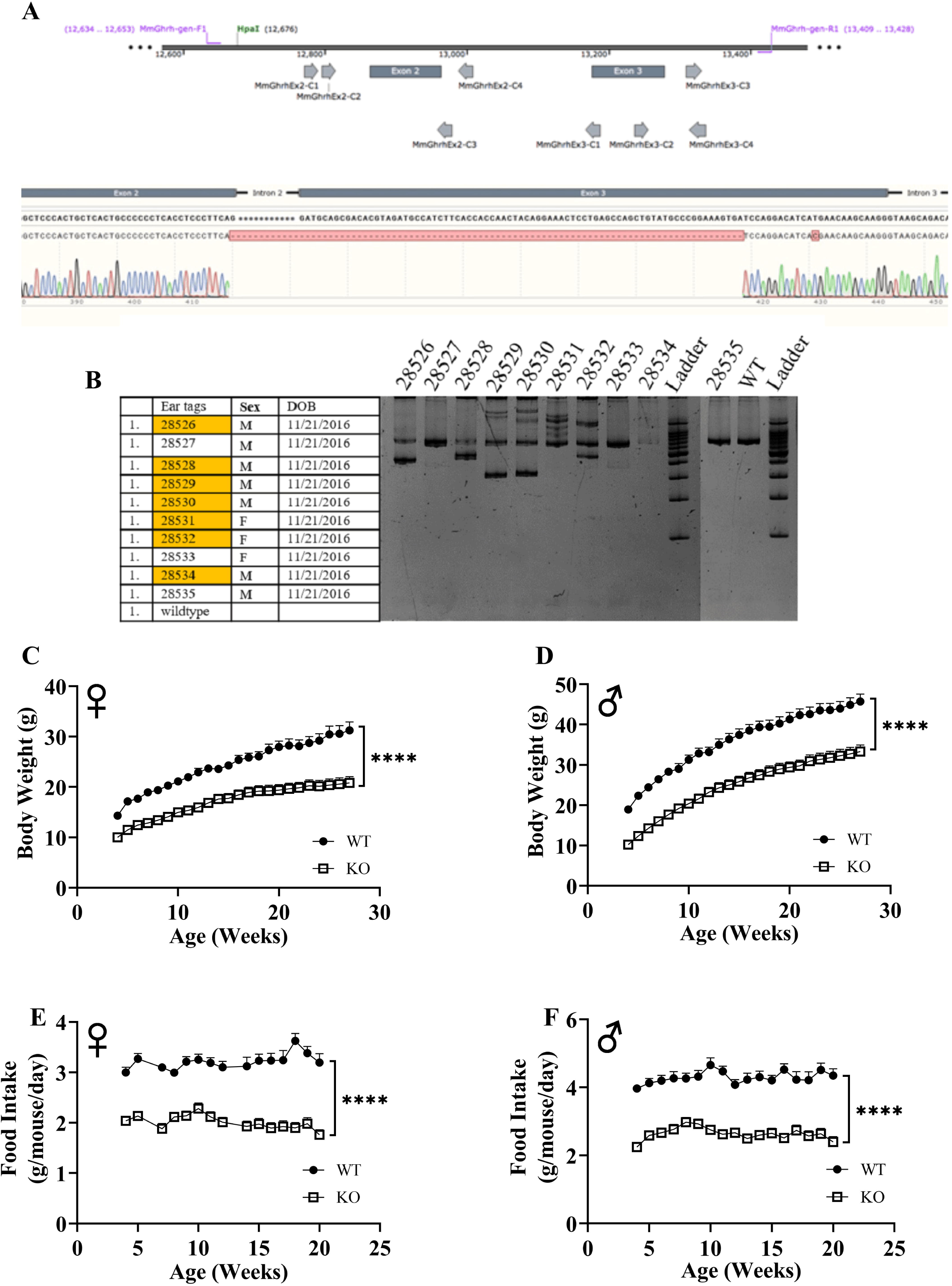
GHRH knockout with CRISPR technology. Location of guide RNAs with respect to exon 2 and exon 3 of GHRH and DNA sequencing chromatogram of mutant GHRH gene between exon 2 and intron 3 (A). Identification of mutations introduced by CRISPR/Cas9 in GHRH gene in founder animals by PCR analysis. 10 G0 pups were tested for indels or deletions. 28528 had a 291 base pairs deletion that eliminates the splice donor site at exon 2, intron 2-3 and a large part of Exon 3 (77 base pairs out of 102 base pairs), showed successful germline transmission (B). Body weights of female (C) and male (D) WT and GHRH^-/-^ mice from weaning to adulthood. Food intake per mice per day of female (E) and male (F) WT and GHRH^-/-^ mice. Female WT n=11, GHRH^-/-^ n=14, male WT n=11, GHRH^-/-^ n=15. Each bar represents mean ± SEM. Statistical analysis was performed by unpaired Student’s t-test with Welch’s correction; ****p<0.0001.

We measured body weight and food consumption of GHRH^-/-^ and WT littermates longitudinally. Male and female GHRH^-/-^ mice were significantly lighter than their littermate controls (Fig. 1C, D). In addition, GHRH^-/-^ mice consumed dramatically fewer calories than WT littermates (Fig. E, F). GHRH is a hypothalamic peptide that controls both the synthesis and secretion of GH [18]. Previously, knockout of GHRH gene was shown to result in significant reduction of GH expression in pituitary and IGF-1 expression in liver [17]. To confirm successful loss of function of GHRH, we measured mRNA levels of GH in pituitary and IGF-1 in liver. We observed that expression of both genes was significantly decreased in both male and female GHRH^-/-^ mice when compared to their WT littermates (Fig. 2A B, E, and F). These results confirm the reduction in GH/IGF-1 signaling. We did not observe significant change in prolactin expression in the pituitary (Fig. 2C, D).

**Figure 2.**
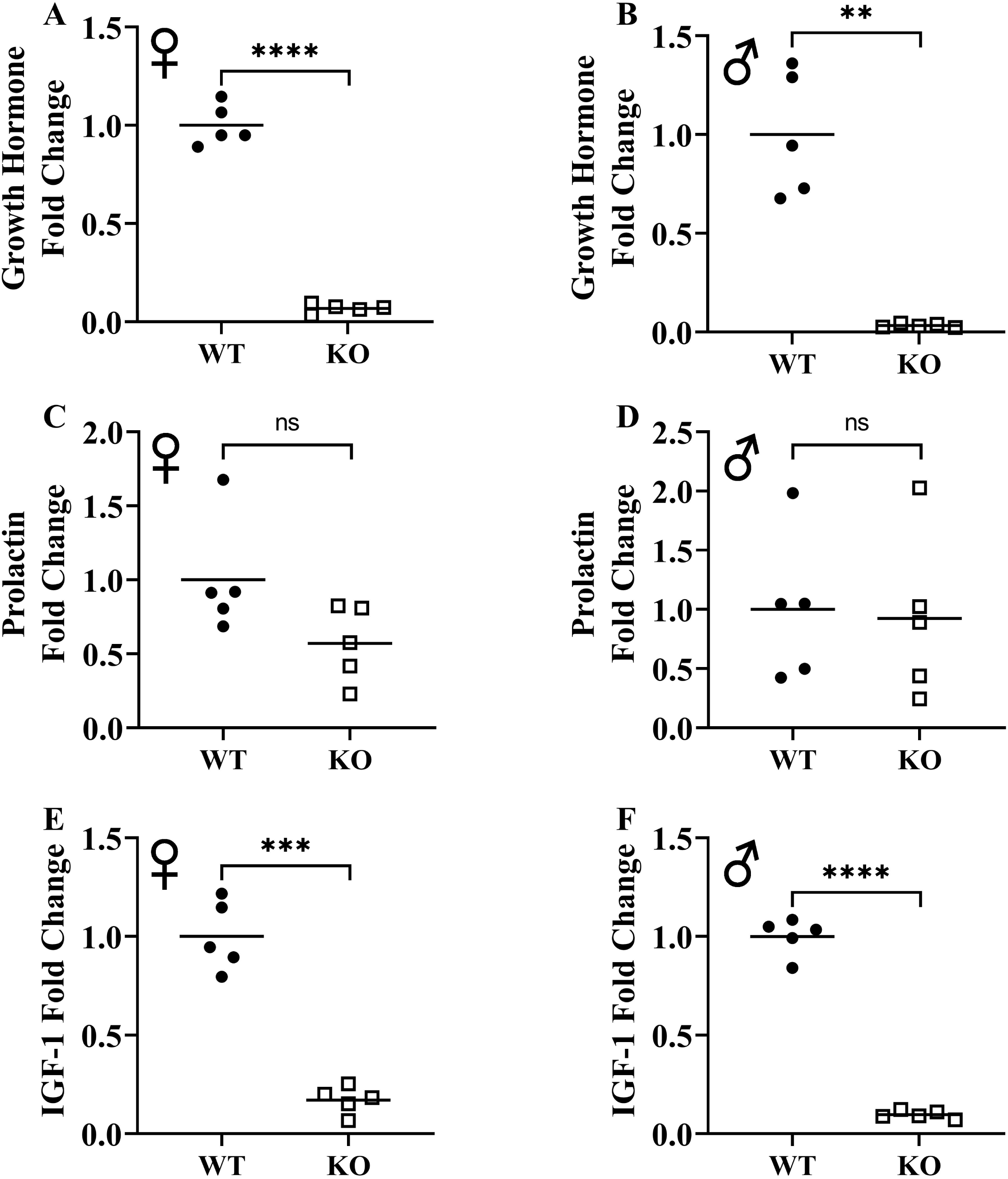
Suppression of growth hormone signaling. Expression of growth hormone gene in pituitary gland in female (A) and male (B) WT and GHRH^-/-^ mice. Expression of prolactin gene in pituitary gland in female (C) and male (D) WT and GHRH^-/-^ mice. Expression of IGF-1 gene in liver in female (E) and male (F) WT and GHRH^-/-^ mice. Expression levels are shown as fold change compared to WT mice. For all biological groups n=5. Each bar represents means. Statistical analysis was performed by unpaired Student’s t-test with Welch’s correction; ns= not significant, **p<0.01, ***p<0.001, ****p<0.0001.

The effect of GH signaling on body weight and composition is well documented in GH related mutant mice [21–23]. Therefore, we performed dual-energy X-ray absorptiometry (DXA) to study the effects of reduced GH signaling on body composition parameters in our knockout model of GHRH. Absolute BMD, BMC, and lean mass values were significantly lower in both male and female GHRH^-/-^ mice compared to WT littermates (Fig. 3C-H). However, to account for the significant body weight differences between GHRH^-/-^ and WT littermates, we used analysis of covariance (ANCOVA) method, which revealed that BMD, BMC, and lean mass were significantly reduced (Fig. 4A-F), whereas, fat mass was significantly increased in GHRH^-/-^ mice (Fig. 4G, H).

**Figure 3.**
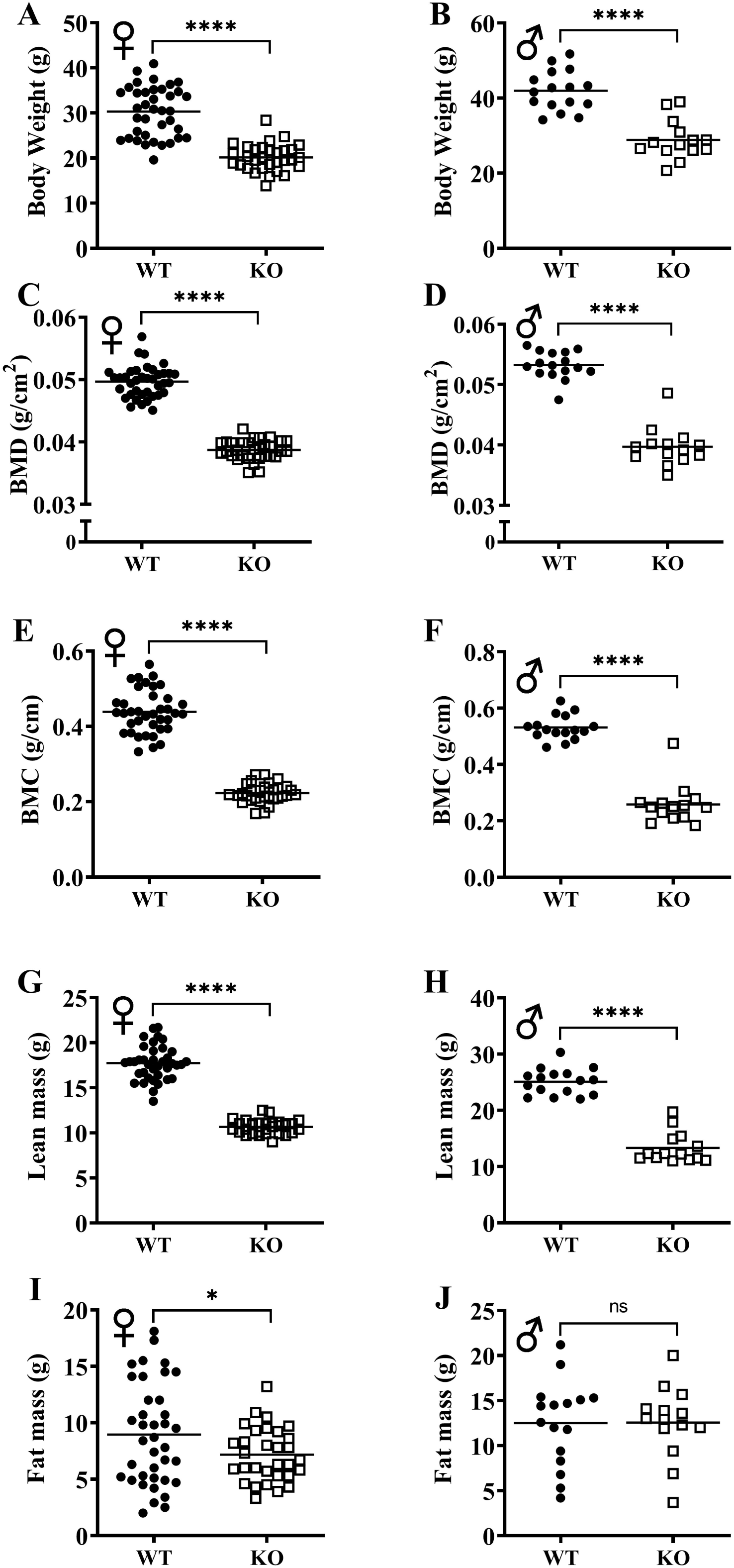
GH-deficiency alters absolute body composition parameters. Body composition parameters: BMD (B, D), BMC (E, F), lean mass (G, H) and fat mass (I, J) were measured by DXA. Female WT n=38, GHRH^-/-^ n=31, male WT n=16, GHRH^-/-^ n=14. Each bar represents mean. Statistical analysis was performed by unpaired Student’s t-test with Welch’s correction; ns= not significant, *p<0.05, ****p<0.0001.

**Figure 4.**
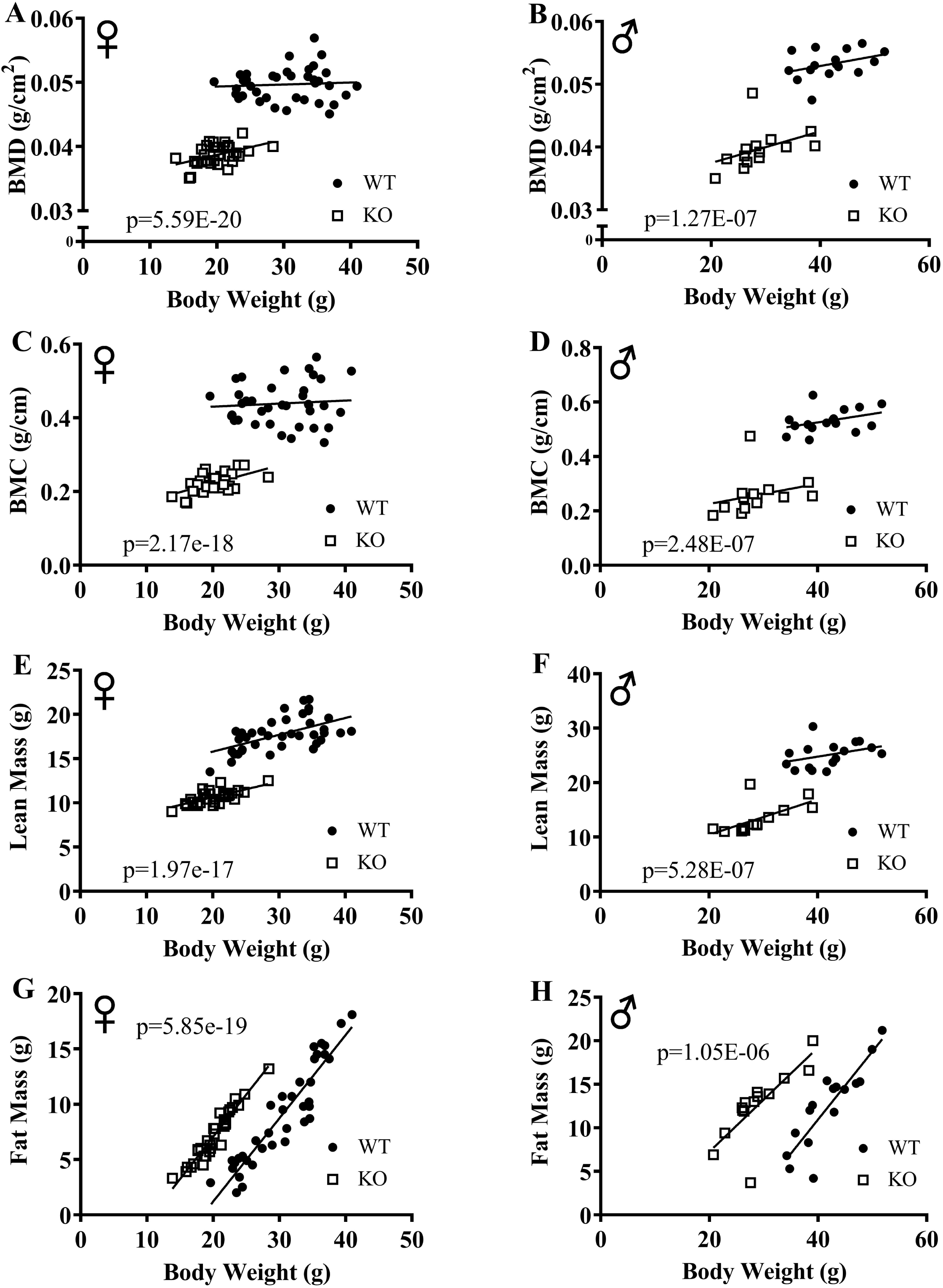
ANCOVA shows GH-deficiency alters body composition parameters. Body composition parameters were measured by DXA. Body composition parameters are plotted on the y-axis and body weights are plotted on the x-axis (A-H). Relationship between body weight and bone mineral density in female (A) and male (B) WT and GHRH^-/-^ mice. Relationship between body weight and bone mineral content in female (C) and male (D) WT and GHRH^-/-^ mice. Relationship between body weight and lean mass in female (E) and male (F) WT and GHRH^-/-^ mice. Relationship between body weight and fat mass in female (G) and male (H) WT and GHRH^-/-^ mice. Female WT n=38, GHRH^-/-^ n=31, male WT n=16, GHRH^-/-^ n=14. The WT and GHRH^-/-^ groups were statistically analyzed with ANCOVA method, which was used to calculate p values, shown on each panel.

To explore the effect of GH deficiency on metabolic phenotype of mice, we measured RER using indirect calorimetry. Fig. 5A and B provides overview of RER values for each hour recorded for 6 days for female and male mice, respectively. Notably, the RER of both male and female GHRH^-/-^ mice decreased much more rapidly during the transition between dark and light cycles compared with WT littermates (Fig. 5A, B). RER measurements collected for 6 days were averaged into a single day (Fig. 5C, D). These results clearly show significantly lower RER for both male and female GHRH^-/-^ mice compared to WT littermates during the light cycle, but not dark cycle (Fig. 5C, D). The comparison of RER measurements collected from 6 light and 6 dark cycles confirm that the significant difference in the metabolism of GHRH^-/-^ mice functions in circadian manner (Fig. 5E-H).

**Figure 5.**
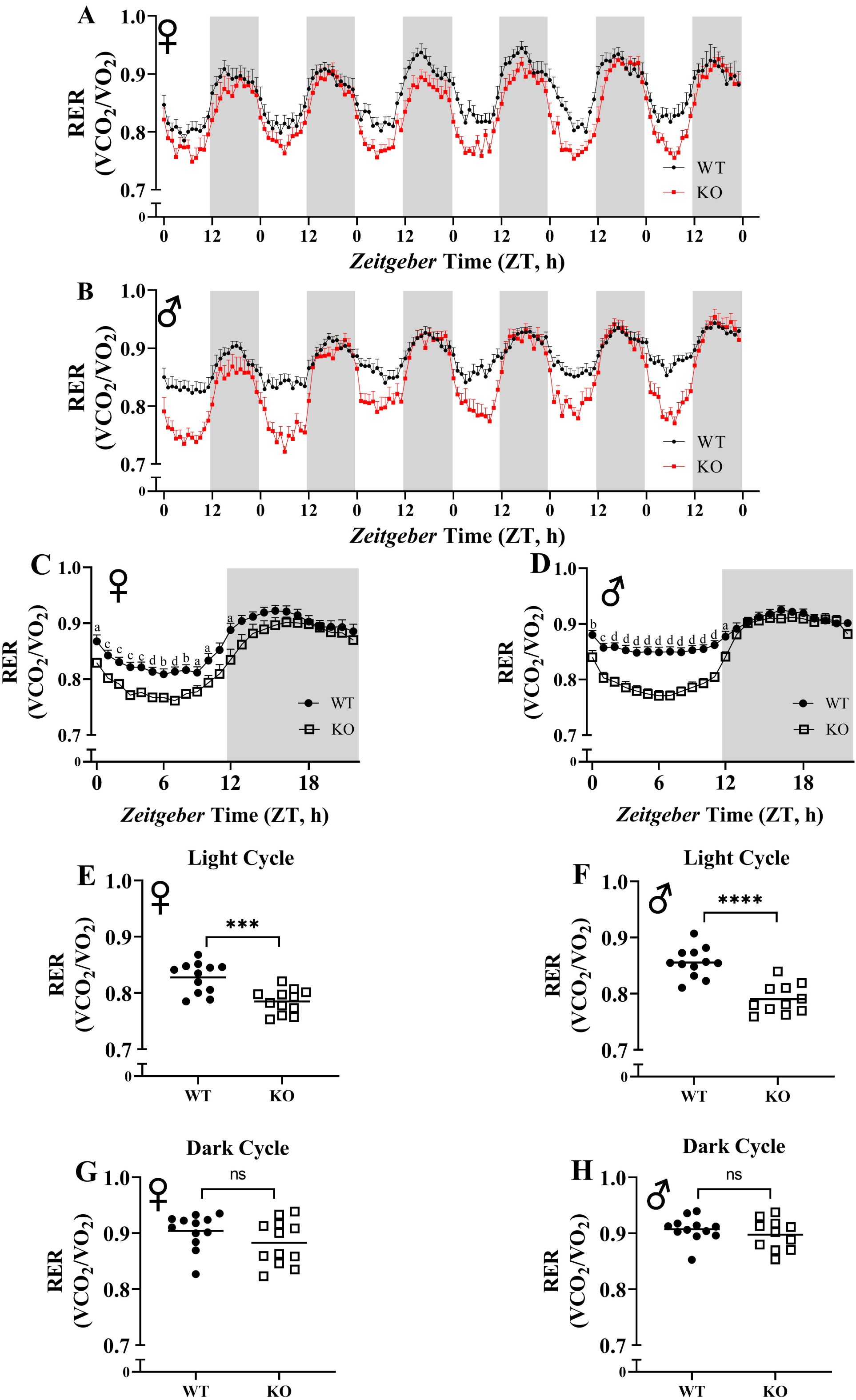
Respiratory exchange ratio (VCO_2_/VO_2_). RER values were calculated by dividing VCO_2_ with VO_2_. 6 days of female (A) and male (B) WT and GHRH^-/-^ mice RER values are shown. Hourly averaged RER values on day of female (C) and male (D) WT and GHRH^-/-^ mice. Overall averaged RER values are shown as light (E, F) and dark cycles (G, H) for female (E, G) and male (F, H) WT and GHRH^-/-^ mice. Female WT n=12, GHRH^-/-^ n=12, male WT n=12, GHRH^-/-^ n=11. Each bar represents mean ± SEM. Statistical analysis was performed by unpaired Student’s t-test with Welch’s correction; ns= not significant, a; *p<0.05, b; **p<0.01, c; ***p<0.001, d; ****p<0.0001.

Fig. 6A, B, E, and F show mean hourly oxygen consumption (VO_2_) and carbon dioxide production (VCO_2_) of male and female mice for 6 days. All mice presented diurnal rhythm of higher VO_2_, VCO_2_ during the dark cycles compared to VO_2_, and VCO_2_ measured during the light cycles. Fig. 5C, D, G, and H show VO_2_ and VCO_2_ measurements collected for 6 days were averaged into a single day. Male and female GHRH^-/-^ have significantly lower absolute VO_2_ and VCO_2_ compared to WT littermates (Fig. 5C, D, G, and H). Overall averages of light and dark cycle absolute VO_2_ and VCO_2_ measurements were significantly lower in GHRH^-/-^ female and male mice compared to WT littermates in both light and dark cycles (Fig. 7A-H). In parallel, ANCOVA, which controls for differences in body weight, showed male and female GHRH^-/-^ mice have significantly lower respiration rates compared to WT littermates, during both light and dark cycles (Fig. 8A-H).

**Figure 6.**
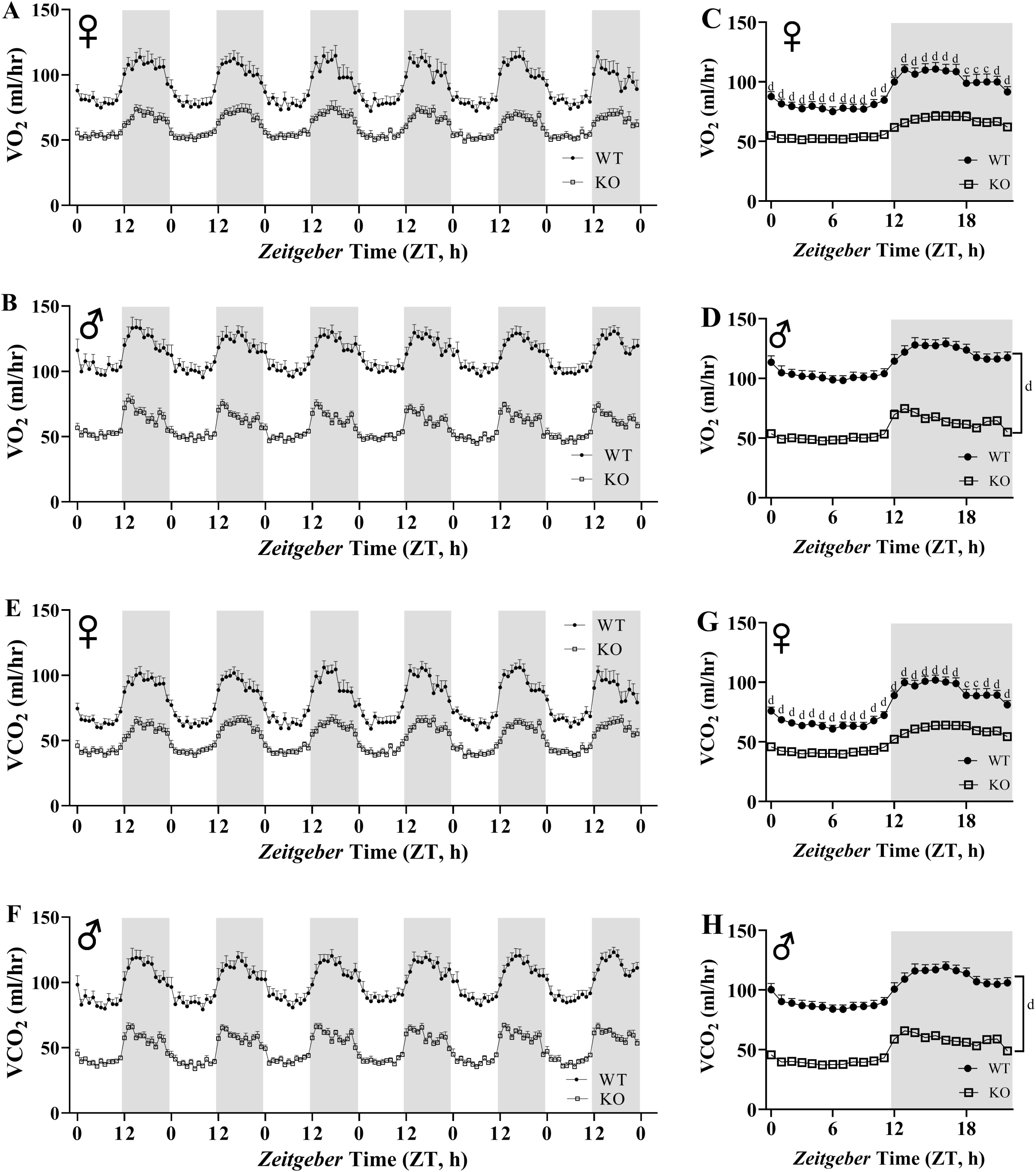
GHRH^-/-^ mice have significantly reduced absolute VO_2_ and VCO_2_. VO_2_ and VCO_2_ were measured by indirect calorimetry. Hourly averages of respiratory parameters measured for 6 days for female (A, E) and male (B, F) WT and GHRH^-/-^ mice. 6 days of VO_2_ and VCO_2_ data were averaged into a single day for female (C, G) and male (D, H) WT and GHRH^-/-^ mice. Female WT n=12, GHRH^-/-^ n=12, male WT n=12, GHRH^-/-^ n=11. Each bar represents mean ± SEM. Statistical analysis was performed by unpaired Student’s t-test with Welch’s correction; ns= not significant, a; p<0.05, b; p<0.01, c; p<0.001, d; p<0.0001.

**Figure 7.**
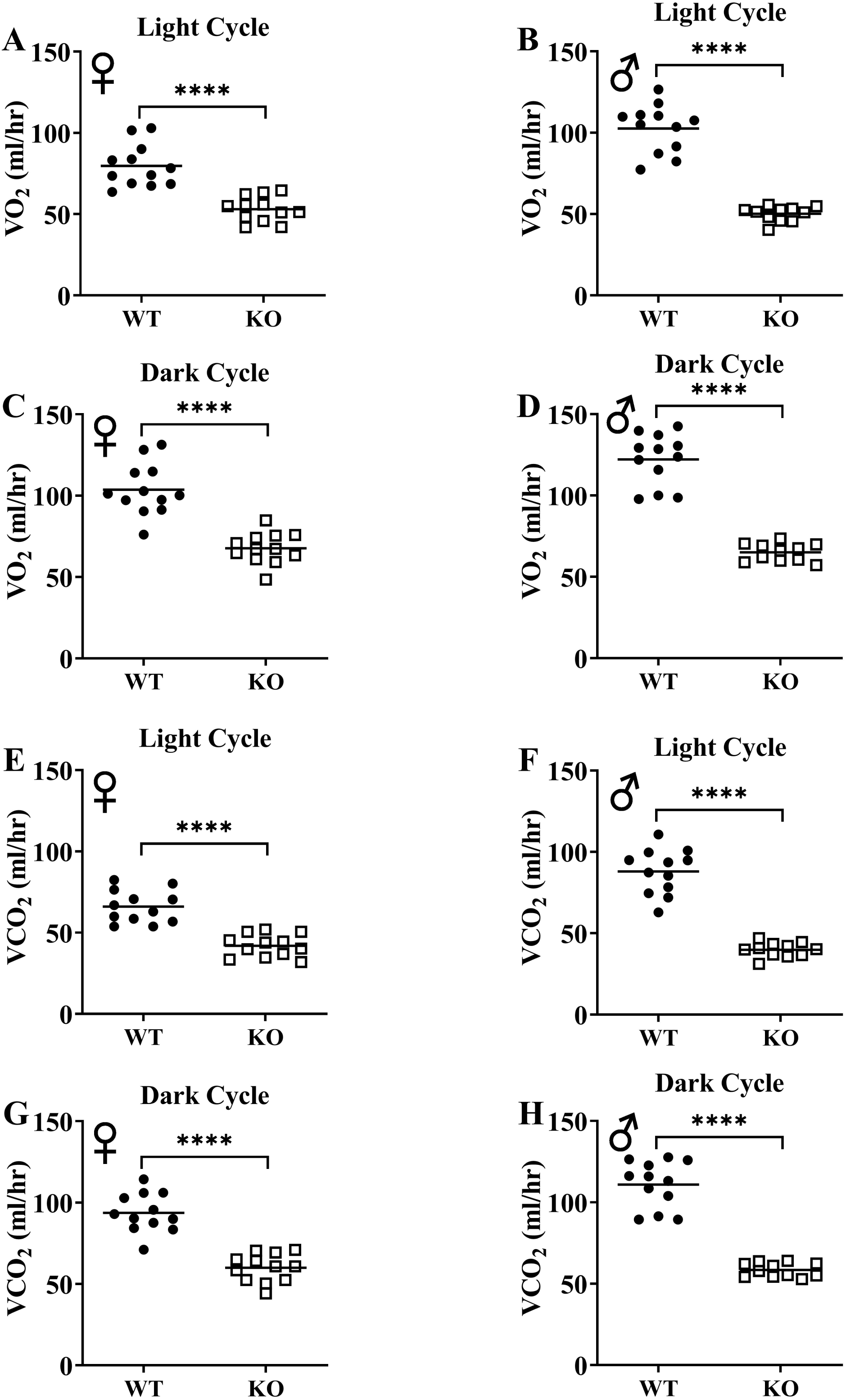
GH-deficiency decreases absolute VO_2_ and VCO_2_. VO_2_ (A-D) and VCO_2_ (E-H) values measured on light and dark cycles were averaged. WT female n=12, KO female n=12, WT male n=12, KO male n=11. Each bar represents mean. Statistical analysis was performed by unpaired Student’s t-test with Welch’s correction; ****p<0.0001.

**Figure 8.**
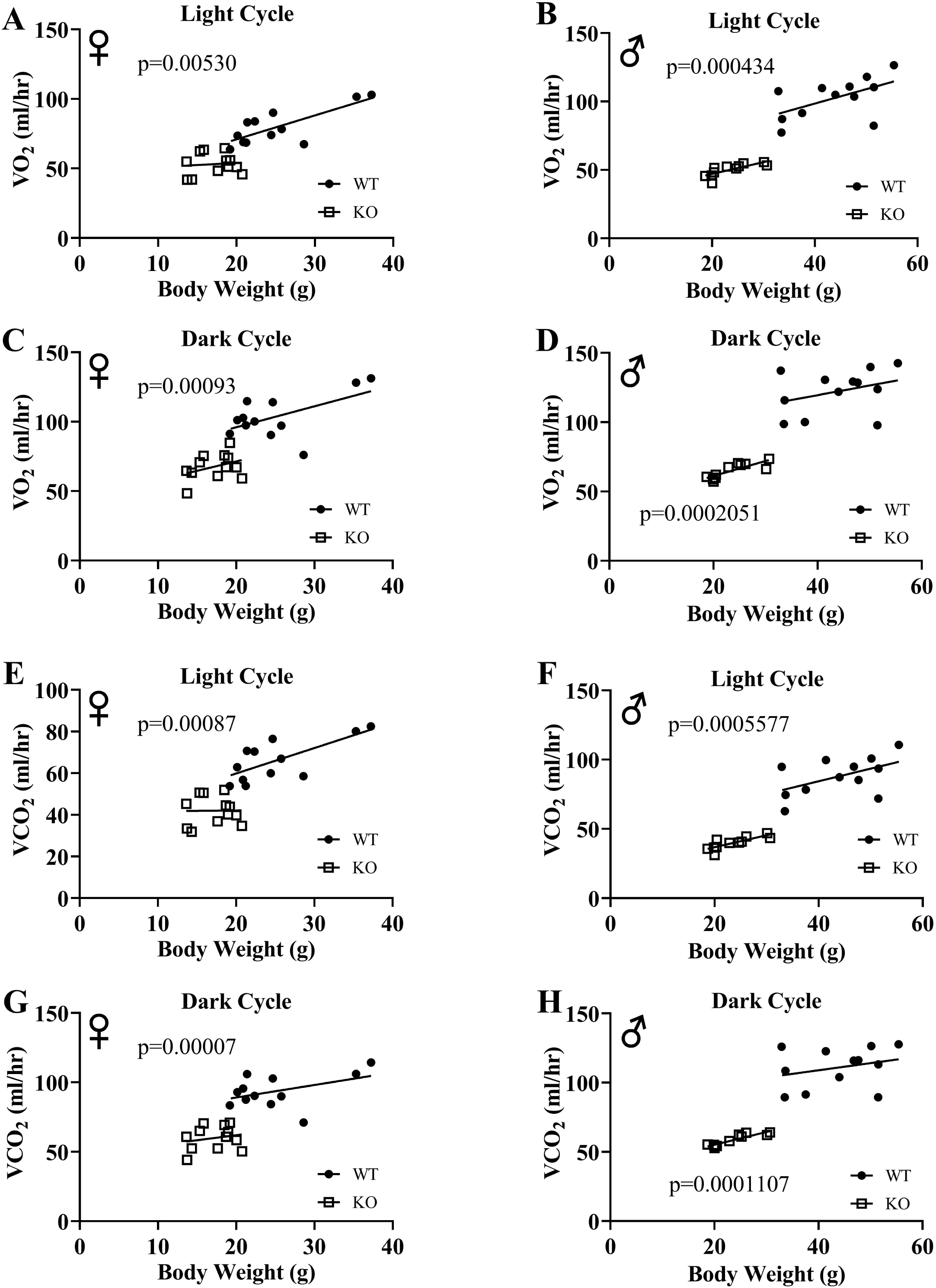
GH-deficiency decreases respiratory parameters. Overall averaged VO_2_ (A-D) and VCO_2_ (E-H) values are plotted on the y-axis and body weights are plotted on the x-axis. Relationship between body weight and VO_2_ in female (A, C) and male (B, D) WT and GHRH^-/-^ mice during light cycle (A, B) and dark cycle (C, D). Relationship between body weight and VCO_2_ in female (E, G) and male (F, H) WT and GHRH^-/-^ mice in light cycles (E, F) and dark cycles (G, H). Female WT n=12, GHRH^-/-^ n=12, male WT n=12, GHRH^-/-^ n=11. The WT and GHRH^-/-^ groups were statistically analyzed with ANCOVA method, which was used to calculate p values, shown on each panel.

We calculated energy expenditure from the respiratory parameters collected by indirect calorimetry to assess the effect of GH-deficiency on metabolic rate. Fig. 9A and B show energy expenditure of female and male, respectively, GHRH^-/-^ mice and WT littermates for 6 days. We averaged this data into a single day showing the dramatic reduction of absolute metabolic rate in both male and female GHRH^-/-^ mice compared to WT littermates (Fig. 9C, D). The analyses of energy expenditure representing the 6 light and 6 dark cycles confirmed the downward shift of metabolic rate in GH-deficient mice (Fig. 10A-D). To control for the effect of significant differences in body weight on energy expenditure, we used ANVOCA, which showed dramatic reduction in metabolic rates of both male and female GHRH^-/-^ mice compared to WT littermates in light and dark cycles (Fig. 9E, H). We measured voluntary physical activity of mice during our indirect calorimetry study. Overall pattern indicates both male and female GHRH^-/-^ mice have reduced activity compared to WT littermates during light and dark cycles (Fig. 11A-D).

**Figure 9.**
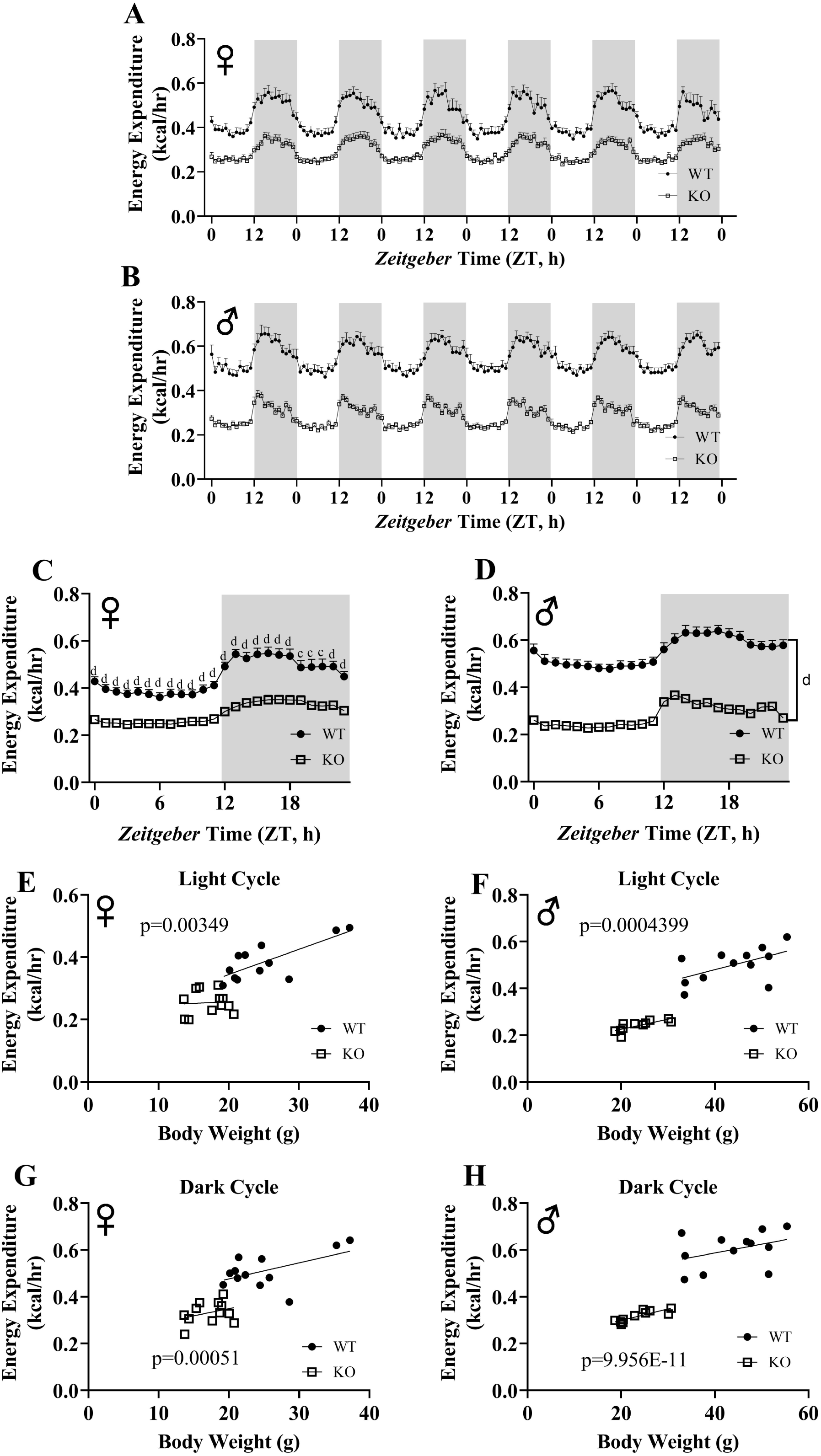
GH-deficiency reduces metabolic rate. Energy expenditure values for 6 days for female (A) and male (B) GHRH^-/-^ and WT mice. 6 days of energy expenditure data were averaged into a single day for female (C) and male (D) mice. Analysis of energy expenditure with body weight as a co-variant for female (E, G) and male (F, H) GHRH^-/-^ and WT mice in light cycles (E, F) and dark cycles (G, H). Female WT n=12, GHRH^-/-^ n=12, male WT n=12, GHRH^-/-^ n=11. Each bar represents mean ± SEM. Statistical analysis was performed by unpaired Student’s t-test with Welch’s correction, c; p<0.001, d; p<0.0001. The WT and GHRH^-/-^ groups were statistically analyzed with ANCOVA method, which was used to calculate p values, shown on panels E-H.

**Figure 10.**
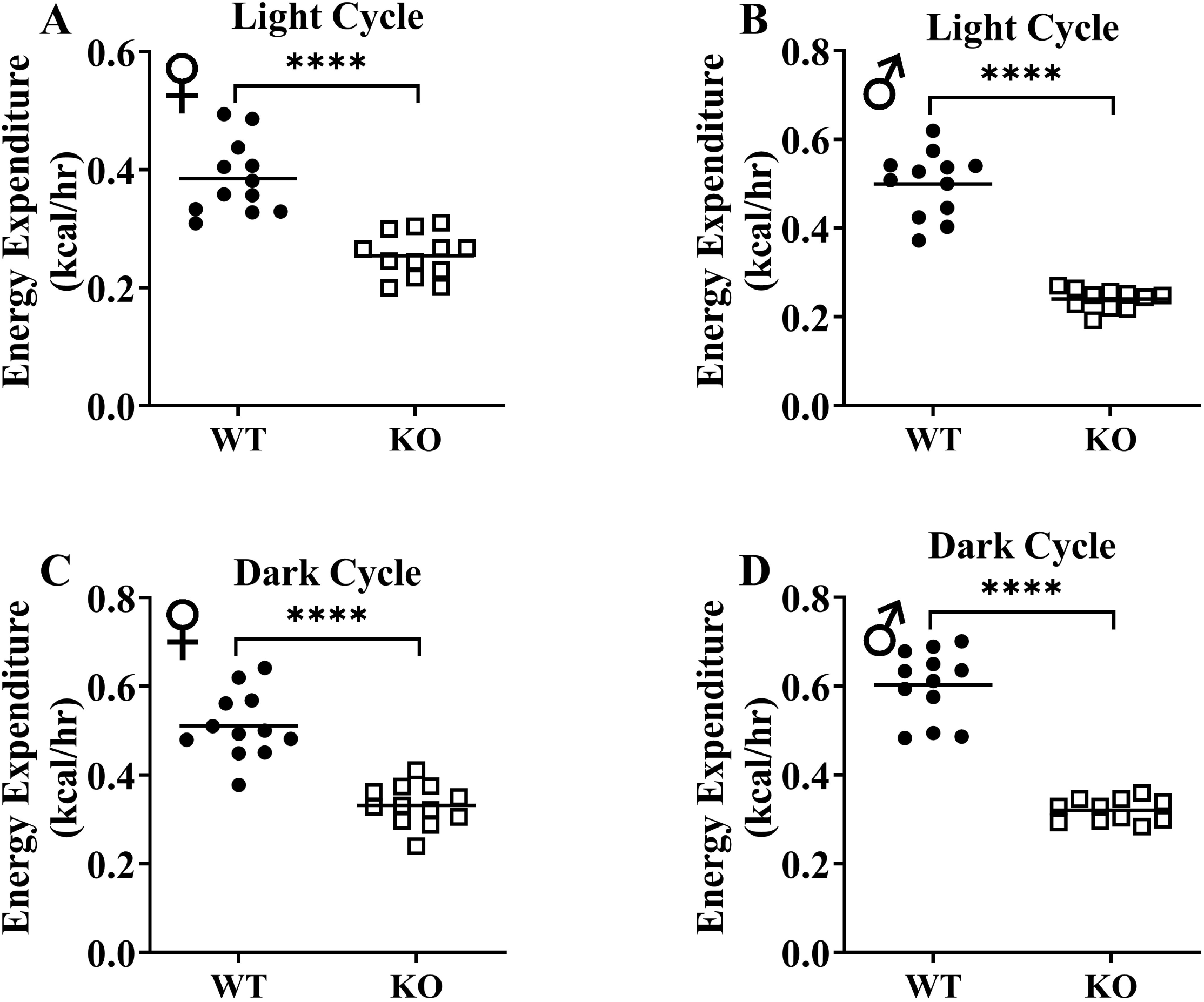
GH-deficiency results in decreased absolute metabolic rate. Overall averaged energy expenditure of WT and GHRH^-/-^ are shown as light (72 hours; A, B) and dark cycles (72 hours; C, D). WT female n=12, KO female n=12, WT male n=12, KO male n=11. Each bar represents mean. Statistical analysis was performed by Student’s t-test with Welch’s correction; ****p<0.0001.

**Figure 11.**
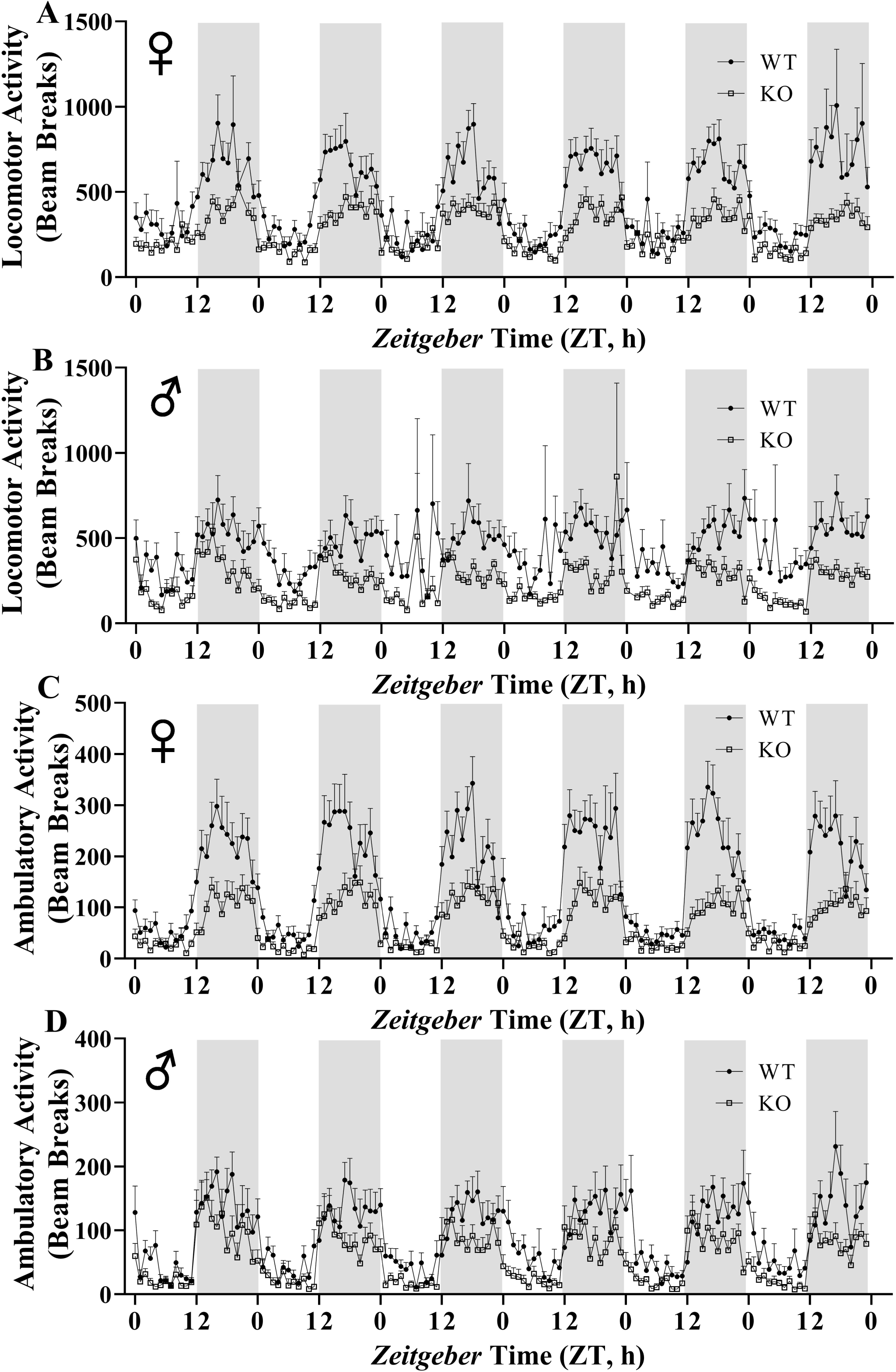
GH-deficiency results in decreased physical activity. Locomotor activity (A, B) and ambulatory activity (C, D) of WT and GHRH^-/-^ mice for 6 days are shown. WT female n=12, KO female n=12, WT male n=12, KO male n=11.

To assess insulin tolerance, we performed intraperitoneal insulin injections and measured blood glucose levels. Upon injection with insulin, glucose concentrations decreased significantly lower levels in GHRH^-/-^ female and male mice than their littermate controls (Fig. 12A, B). Area under the curve (AUC) data were significantly lower GHRH^-/-^ female and male mice than their littermate controls (Fig. 12C, D). To evaluate the glucose homeostasis *in vivo*, we performed intraperitoneal glucose tolerance test (IPGTT). We did not observe any significant differences in blood glucose levels throughout the 2-hour period following glucose injection (Fig. 12E, F). AUC analyses did not reveal any statistical significance due to loss of GHRH (Fig. 12G, H). This data strongly supports the notion that GH deficiency improves insulin sensitivity *in vivo*.

**Figure 12.**
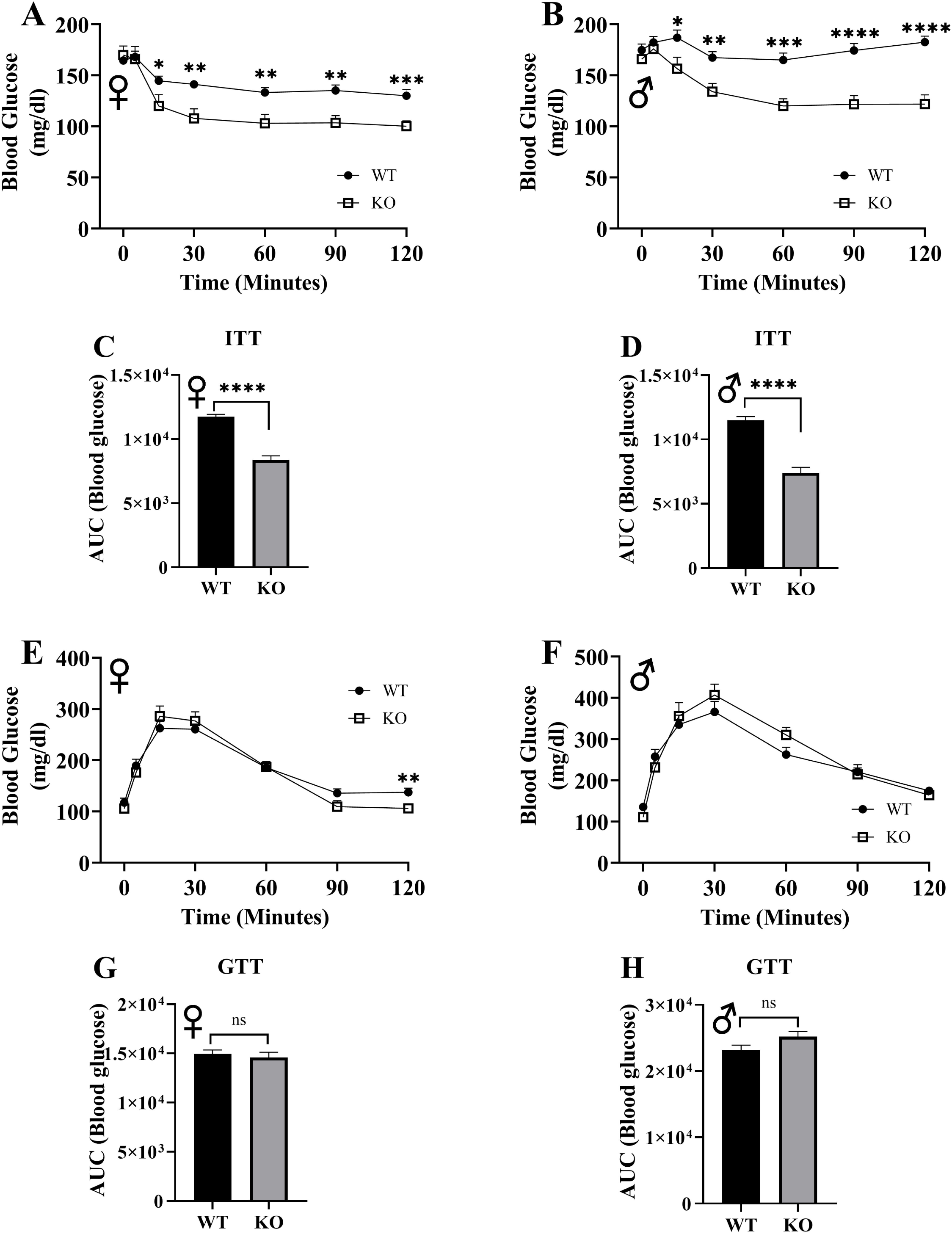
Insulin and glucose tolerance tests. GHRH^-/-^ and WT mice were fasted for 4 hours and injected with 1 IU porcine insulin per kg of body weight. Blood glucose levels of female (A) and male (B) were measured the following 2 hours. Area under the curve analyses for female (C) and male (D) mice are shown. GHRH^-/-^ and WT mice were fasted overnight and injected with 1 g glucose per kg of body weight. Blood glucose levels of female (E) and male (F) were measured the following 2 hours. Area under the curve analyses for female (G) and male (H) mice are shown. Female WT n=14, GHRH^-/-^ n=13, male WT n=14, GHRH^-/-^ n=13-14. Each bar represents mean ± SEM. Statistical analysis was performed by unpaired Student’s t-test with Welch’s correction; *p<0.05, **p<0.01, ***p<0.001, ****p<0.0001.

## Discussion

Mouse GHRH gene has 5 exons and is located on chromosome 2. Exon 1 does not include protein coding sequences, exon 2 and a portion of exon 3 encodes for the signal peptide. Most of exon 3 and a portion of exon 4 encode for the mature GHRH peptide [24, 25]. Previously, GHRH gene was knocked out in mice using a neomycin resistance cassette to replace parts of both exon 2 and 3 [17]. This approach introduces the possibility of passenger flanking alleles/mutations, which is one explanation for phenotypic variations between mice generated with CRISPR/Cas9-based gene-editing and classical knockout method [26, 27]. Agouti gene and GHRH^-/-^ alleles co-segregate in the mouse model generated by Salvatori lab [17]. Previously, GHRH^-/-^ mice were shown to have increased adiposity and insulin sensitivity, which are the two key parameters linked to longevity in different knockout models of the growth hormone pathway. However, insulin resistance and obesity are both linked to agouti gene expression [20]. Therefore, we generated a ‘clean’ model for GHRH^-/-^ mice using the CRISPR/Cas9 system, which provides precise genome-editing without leaving any exogenous DNA sequences behind, eliminating the possibility of passenger alleles/mutations influencing the phenotype resulting from the GH-deficiency [28]. It has been shown in the literature that CRISPR/Cas9 method can result in off-target mutations. We have outcrossed our mice several generations to minimize the possibility of off-target mutations in the genome. Some of the phenotypic variability observed in transgenic animals has been attributed to the genetic background of animal models [29]. To rule out this possibility, we used GHRH^-/-^ mice on mixed genetic background for this study.

Indirect calorimetry provides highly sensitive, accurate, and noninvasive measurements of energy expenditure and substrate utilization of live animals [30]. One important technical issue for indirect calorimetry is the duration of acclimation period. Generally, data collection by indirect calorimetry is limited to 24 hours, which usually takes place after a 24 hours acclimation period [5, 31, 32]. Our mice are housed as a group of seven in each cage with room temperature between 20–23°C. Temperature within the cage is thought to be closer to the thermoneutral zone, where energy expenditure required for maintaining body temperature is at its lowest. Moving mice from group housing to single housing is expected to increase their expenditure [33]. For this reason, it is critical for mice to adjust to cold stress before any respiratory measurements are performed. In order to obtain accurate and reliable measurements from indirect calorimetry, we acclimated mice in the respiratory chambers for 7 days.

Interpretation of metabolic and physiological parameters has been problematic due to differences in body weight in some genetic models [34]. In order to compensate for these differences, researchers have argued for different methods of analysis, including ratio-based normalization and allometric scaling [35]. Studies illustrated that these methods are improper and result in flawed conclusions [34–37]. Analysis of covariance (ANCOVA) has been promoted as the suitable method of analysis for physiological parameters that are influenced by variables such as body weight [36, 38, 39]. We utilized this unbiased statistical approach to control for the influence of body weight on body composition and indirect calorimetry data.

The inverse relationship between lifespan and body size within species has been observed not only in mice but also, in rats, dogs, horses and humans [6, 40–42]. Decreased body size is one of the strongest phenotypic characteristics of growth hormone deficiency models [5, 8]. Analyses of body composition parameters with DXA revealed remarkable effects of reduced GH signaling on bone, lean, and fat tissues. Absolute BMD, BMC, lean mass measurements are significantly lower in GHRH^-/-^ female and male mice compared to littermate controls. Reduced absolute lean mass and BMD have been documented in the GHR^-/-^ model [23]. In addition, Ames dwarf mice have significantly reduced absolute lean mass and BMC [43]. For BMC, BMD, and lean mass, comparison of absolute measurements and ANCOVA using body weight as a co-variant resulted in exact same conclusions. We did not observe great differences in absolute fat mass between GHRH^-/-^ female and male mice and WT littermates. However, ANCOVA demonstrated that fat mass adjusted for body weight is significantly increased in mice lacking GHRH.

We aimed to utilize indirect calorimetry to investigate the metabolic effects, which are associated with reduced GH signaling and may be related to improved longevity. Analyses of absolute energy expenditure and body weight adjusted energy expenditure values lead to the same conclusion that our novel GHRH^-/-^ mice have significantly decreased metabolic rates compared to WT littermates. Another study using Ames dwarf and GHR^-/-^ found significantly decreased absolute energy expenditures for both models [31]. Using indirect calorimetry, we examined the RER, which is a unitless ratio obtained by dividing VCO_2_ by VO_2_. RER is close to 0.7 when mice metabolize fat as an energy source and RER is close to 1.0 when mice metabolize carbohydrates [44]. Our results demonstrate that loss of GH-deficiency is significantly associated with lower RER during the light cycle, but not during the dark cycle in both female and male mice. This suggests that GHRH^-/-^ mice have a greater level of fat utilization, which appears to occur in a circadian pattern. Previous studies have shown lower RER for Ames and GHR^-/-^ mice during both light and dark cycles [31]. However, these animals were only acclimated for 24 hours and data were collected in 24 hours. We acclimate our experimental animals for 7 days and collect data for 6 days. This technical difference is most likely to be the reason for the difference in results.

Our study shows that GHRH^-/-^ mice have decreased physical activity compared to their WT littermates. Another study showed increased physical activity in GHRH deficient mice [45]. This study recorded physical activity for only 10 minutes. However, our measurements lasted 6 days. This difference in methodology might be the likely contributor to the difference in results.

In humans, insulin resistance is one of the hallmarks of aging and is associated with chronic conditions such as diabetes, cancer, and cardiovascular disease [46, 47]. One of the physiological characteristics of long-lived GH-related mutant mouse models is elimination of insulin resistance [5, 48, 49]. In this study, our novel CRISPR/Cas9 GHRH^-/-^ female and male mice exhibited insulin sensitivity compared to their WT littermates, affirming the relationship between growth hormone deficiency and increased sensitivity to insulin.

In summary, our study validates our previous findings and establishes the GHRH^-/-^ mouse as an important animal model to study mechanisms of extended longevity and slow aging in mammals. This work provides a practical example to adopt CRISPR/Cas9 targeting system to improve locus specificity and ease of multiplexed targeting in mammalian aging and longevity studies.

## Materials and Methods

### CRISPR/sgRNA design and synthesis

Owing to the small size of the mouse GHRH exons (17.16 kb with 5 exons), eight CRISPR targets with high scores were identified in the sequences flanking exons 2 and 3, including the intronic sequences, using Benchling (www.benchling.com) with a goal to create deletion alleles spanning exons 2 and 3. Single guide RNA (sgRNA) molecules were generated using a cloning-free method as described earlier [50]. Cas9 protein was obtained from MacroLabs at UC Berkeley.

### Generation of G0 (founder) animals and germline transmission of mutant alleles

All animal procedures were performed in accordance with the recommendations in the guide for the care and use of laboratory animals published by the National Institutes of Health. The protocols used were approved and conducted according to the University of Alabama at Birmingham institutional animal care and use committee. Pronuclear injections into C57BL/6J zygotes were performed with a solution of sgRNAs (50 ng/μl each) and Cas9 protein (50 ng/μl per guide). Injected zygotes were implanted into pseudo-pregnant CD1 recipients. Genomic DNA obtained from tail biopsies of putative founder (G_0_) animals were assessed for the presence of mutations in the targeted genes. G_0_ animals were bred to WT C57BL/6J mice for germline transmission of mutant alleles.

### Detecting the presence of indels

Genomic DNA from mouse-tail biopsies was obtained by digesting in lysis buffer (50 mM Tris-HCl pH 8.0, 100 mM EDTA pH 8.0, 100 mM NaCl, 1% SDS) with proteinase K (0.3 mg/ml), followed by a phenol:chloroform extraction and ethanol precipitation procedure. PCRs were set up using the oligonucleotide primers MmGhrh-gen-F1: 5’-CTTGCTTCTCTCACACTTGC-3’; MmGhrh-gen-R1: 5’-TTAAAGGGTCGGAGCAGTAG-3’ with NEB Taq 2x Master Mix. The amplicons (795 base pairs) were subjected to denaturation-slow renaturation to facilitate formation of hetero-duplexes using a thermocycler. These samples were then resolved on polyacrylamide gels (6%) and the resulting mobility profiles used to infer efficiency of CRISPR-Cas9 nuclease activity. Indels or deletions were detected by heteroduplex mobility analysis (HMA) from tail genomic DNA of potential founder animals. PCR amplicons were cloned using the TOPO-TA cloning kit (ThermoFisher/Invitrogen, Carlsbad CA). Colonies were picked from each plate and grown in 1.5 ml liquid cultures to isolate plasmid DNA using an alkaline lysis procedure. Plasmid DNA was sequenced using M13 forward or reverse primers.

### Breeding

Mice were maintained in a fixed light−dark cycle with ad libitum access to food and water. A GHRH heterozygous (^+/-^) G_0_ mouse obtained from the University of Alabama at Birmingham (UAB) genomics core was crossed with a BALB/cByJ for increased genetic diversity, increased fecundity, and reduced aggression. Mice heterozygous for our GHRH mutation were then crossed with littermates to obtain homozygous wild type and homozygous mutant animals. Due to the low number of homozygous mutant mice per litter, litter mate homozygous wild type mice were then breed as controls and homozygous mutant males were breed to littermate female heterozygous females in order to obtain experimental animals.

### Dual-energy X-ray absorptiometry (DXA)

The mice were scanned using the GE Lunar PIXImus DXA with software version 1.45. The mice were anesthetized using an Isoflurane (3%) and oxygen (500ml/min) mixture, delivered by a Surgivet anesthesia machine, and then placed in a prostrate position on the DXA imaging plate and scanned. During the scan, the mice remained anesthetized. For all scans, the head was excluded from the analysis and the data obtained included BMC, BMD, lean mass and fat mass.

### Indirect Calorimetry

Indirect calorimetry was performed using comprehensive lab animal monitoring system (Oxymax-CLAMS; Columbus Instruments Co., Columbus, OH). This system uses zirconia and infrared sensors to monitor oxygen (O_2_) and carbon dioxide (CO_2_), respectively. We performed indirect calorimetry with 24 mice (12 wild type and 12 GHRH^-/-^). We divided mice into 3 groups and collected measurements on 8 animals at a time (4 WT and 4 GHRH^-/-^). The mice were housed in separate respiratory chambers for 7 days for acclimatization before starting the measurements. After a 7-day acclimation period, respiratory parameters of mice were recorded for 6 days with ad-libitum access to standard chow and water. Respiratory samples were measured every 9 minutes per mouse, and the data were averaged for each hour. RER was calculated by dividing VCO_2_ by VO_2_. Energy expenditure was calculated by the equation as energy expenditure = (3.815 + 1.232 × VCO_2_/VO_2_) × VO_2_ [44]. We used infrared beam system in X, Y, and Z coordinates to record physical activity of mice. If the mouse is standing still and starts a repetitive act, such as grooming, it will continuously break the same beam indicating locomotor activity. When the currently broken beam is different from the previous one, activity is counted as ambulatory.

### Glucose and Insulin Tolerance Tests

Overnight-fasted mice underwent glucose tolerance test by intraperitoneal injection with 1 g of glucose per kg of body weight. Blood glucose levels were measured at 0, 5, 15, 30, 60, 90, and 120 minutes. 4-hours fasted mice underwent insulin tolerance test by intraperitoneal injection with 1 IU porcine insulin (Sigma-Aldrich, St. Louis, MO) per kg of body weight. Blood glucose levels were measured at 0, 5, 15, 30, 60, 90, and 120 minutes.

### Real-Time Quantitative PCR

RNA was harvested from tissues using RNeasy plus kit (Qiagen, Hilden, Germany). Total RNA was reverse transcribed with LunaScript RT SuperMix Kit (New England Biolabs, Ipswich, MA). Real-time quantitative PCR was performed using a QuantStudio 3 with a PowerUp SYBR green master mix (ThermoFisher Scientific, Waltham, MA). Glyceraldehyde-3-phosphate dehydrogenase (GAPDH) or beta-actin expression was used to normalize gene of interest in each sample. Real-time quantitative PCRs were set up using the oligonucleotide primers Mm GAPDH F1 5-CCTGGAGAAACCTGCCAAGTATGATG-3’; Mm GAPDH R1 5-AAGAGTGGGAGTTGCTGTTGAAGTC-3’, Mm Actb F4 5’-TCTTTGCAGCTCCTTCGTTGCC-3; Mm Actb R4 5’-CTGACCCATTCCCACCATCACAC-3’, Mm IGF-1 F2 5’-CATAGTACCCACTCTGACCTGCTGTG-3’; Mm IGF-1 R2 5’-CGCCAGGTAGAAGAGGTGTGAAGAC-3’; Mm GH F1 5’-TGGCTACAGACTCTCGG-3’; Mm GH R1 5’-AGAGCAGGCAGAGCAGGCTGA-3’. Fold change was obtained by calculating 2^−ΔΔCt^.

### Statistical Analyses

The unpaired Student’s t-test with Welch’s correction was used for statistical analysis. Statistical significance was established at p<0.05, two-tailed. We used (generalized linear model) GLM package with R software for analysis of indirect calorimetry and body composition data. When interaction was not found, the code was run without the interaction term. Our GLM models were validated with the MMPC’s (National Mouse Metabolic Phenotyping Center, https://www.mmpc.org/shared/regression.aspx) energy expenditure analysis tool. Graphs were generated with GraphPad Prism 8 (San Diego, CA).

## Acknowledgements

We thank Sofia Canlas, Joseph Jablonsky, Whitney Turner, and Matthew Joyner for technical assistance. We also thank other members of the Sun lab for their helpful discussion and comments on the revision of the manuscript.

## References

1. Lopez-Otin C, Blasco MA, Partridge L, Serrano M, Kroemer G. The hallmarks of aging. Cell. 2013; 153: 1194–217.

2. Kenyon C, Chang J, Gensch E, Rudner A, Tabtiang R. A C. elegans mutant that lives twice as long as wild type. Nature. 1993; 366: 461–4.

3. Flatt T, Min KJ, D’Alterio C, Villa-Cuesta E, Cumbers J, Lehmann R, Jones DL, Tatar M. Drosophila germ-line modulation of insulin signaling and lifespan. Proc Natl Acad Sci U S A. 2008; 105: 6368–73.

4. Coschigano KT, Holland AN, Riders ME, List EO, Flyvbjerg A, Kopchick JJ. Deletion, but not antagonism, of the mouse growth hormone receptor results in severely decreased body weights, insulin, and insulin-like growth factor I levels and increased life span. Endocrinology. 2003; 144: 3799–810.

5. Sun LY, Spong A, Swindell WR, Fang Y, Hill C, Huber JA, Boehm JD, Westbrook R, Salvatori R, Bartke A. Growth hormone-releasing hormone disruption extends lifespan and regulates response to caloric restriction in mice. Elife. 2013; 2: e01098.

6. Flurkey K, Papaconstantinou J, Miller RA, Harrison DE. Lifespan extension and delayed immune and collagen aging in mutant mice with defects in growth hormone production. Proc Natl Acad Sci U S A. 2001; 98: 6736–41.

7. Brown-Borg HM, Borg KE, Meliska CJ, Bartke A. Dwarf mice and the ageing process. Nature. 1996; 384: 33.

8. Coschigano KT, Clemmons D, Bellush LL, Kopchick JJ. Assessment of growth parameters and life span of GHR/BP gene-disrupted mice. Endocrinology. 2000; 141: 2608–13.

9. Bartke A, Sun LY, Longo V. Somatotropic signaling: trade-offs between growth, reproductive development, and longevity. Physiol Rev. 2013; 93: 571–98.

10. van der Spoel E, Jansen SW, Akintola AA, Ballieux BE, Cobbaert CM, Slagboom PE, Blauw GJ, Westendorp RGJ, Pijl H, Roelfsema F, van Heemst D. Growth hormone secretion is diminished and tightly controlled in humans enriched for familial longevity. Aging Cell. 2016; 15: 1126–31.

11. Sornson MW, Wu W, Dasen JS, Flynn SE, Norman DJ, O’Connell SM, Gukovsky I, Carriere C, Ryan AK, Miller AP, Zuo L, Gleiberman AS, Andersen B, et al. Pituitary lineage determination by the Prophet of Pit-1 homeodomain factor defective in Ames dwarfism. Nature. 1996; 384: 327–33.

12. Li S, Crenshaw EB, 3rd, Rawson EJ, Simmons DM, Swanson LW, Rosenfeld MG. Dwarf locus mutants lacking three pituitary cell types result from mutations in the POU-domain gene pit-1. Nature. 1990; 347: 528–33.

13. Ikeno Y, Bronson RT, Hubbard GB, Lee S, Bartke A. Delayed occurrence of fatal neoplastic diseases in ames dwarf mice: correlation to extended longevity. J Gerontol A Biol Sci Med Sci. 2003; 58: 291–6.

14. Chandrashekar V, Bartke A, Coschigano KT, Kopchick JJ. Pituitary and testicular function in growth hormone receptor gene knockout mice. Endocrinology. 1999; 140: 1082–8.

15. Hauck SJ, Hunter WS, Danilovich N, Kopchick JJ, Bartke A. Reduced levels of thyroid hormones, insulin, and glucose, and lower body core temperature in the growth hormone receptor/binding protein knockout mouse. Exp Biol Med (Maywood). 2001; 226: 552–8.

16. Zhou Y, Xu BC, Maheshwari HG, He L, Reed M, Lozykowski M, Okada S, Cataldo L, Coschigamo K, Wagner TE, Baumann G, Kopchick JJ. A mammalian model for Laron syndrome produced by targeted disruption of the mouse growth hormone receptor/binding protein gene (the Laron mouse). Proc Natl Acad Sci U S A. 1997; 94: 13215–20.

17. Alba M, Salvatori R. A mouse with targeted ablation of the growth hormone-releasing hormone gene: a new model of isolated growth hormone deficiency. Endocrinology. 2004; 145: 4134–43.

18. Muller EE, Locatelli V, Cocchi D. Neuroendocrine control of growth hormone secretion. Physiol Rev. 1999; 79: 511–607.

19. Smith SR, Gawronska-Kozak B, Janderova L, Nguyen T, Murrell A, Stephens JM, Mynatt RL. Agouti expression in human adipose tissue: functional consequences and increased expression in type 2 diabetes. Diabetes. 2003; 52: 2914–22.

20. Klebig ML, Wilkinson JE, Geisler JG, Woychik RP. Ectopic expression of the agouti gene in transgenic mice causes obesity, features of type II diabetes, and yellow fur. Proc Natl Acad Sci U S A. 1995; 92: 4728–32.

21. List EO, Sackmann-Sala L, Berryman DE, Funk K, Kelder B, Gosney ES, Okada S, Ding J, Cruz-Topete D, Kopchick JJ. Endocrine parameters and phenotypes of the growth hormone receptor gene disrupted (GHR-/-) mouse. Endocr Rev. 2011; 32: 356–86.

22. Bonkowski MS, Pamenter RW, Rocha JS, Masternak MM, Panici JA, Bartke A. Long-lived growth hormone receptor knockout mice show a delay in age-related changes of body composition and bone characteristics. J Gerontol A Biol Sci Med Sci. 2006; 61: 562–7.

23. Berryman DE, List EO, Palmer AJ, Chung MY, Wright-Piekarski J, Lubbers E, O’Connor P, Okada S, Kopchick JJ. Two-year body composition analyses of long-lived GHR null mice. J Gerontol A Biol Sci Med Sci. 2010; 65: 31–40.

24. Mayo KE, Cerelli GM, Lebo RV, Bruce BD, Rosenfeld MG, Evans RM. Gene encoding human growth hormone-releasing factor precursor: structure, sequence, and chromosomal assignment. Proc Natl Acad Sci U S A. 1985; 82: 63–7.

25. Frohman MA, Downs TR, Chomczynski P, Frohman LA. Cloning and characterization of mouse growth hormone-releasing hormone (GRH) complementary DNA: increased GRH messenger RNA levels in the growth hormone-deficient lit/lit mouse. Mol Endocrinol. 1989; 3: 1529–36.

26. Lee JH, Park JH, Nam TW, Seo SM, Kim JY, Lee HK, Han JH, Park SY, Choi YK, Lee HW. Differences between immunodeficient mice generated by classical gene targeting and CRISPR/Cas9-mediated gene knockout. Transgenic Res. 2018; 27: 241–51.

27. Szabo R, Samson AL, Lawrence DA, Medcalf RL, Bugge TH. Passenger mutations and aberrant gene expression in congenic tissue plasminogen activator-deficient mouse strains. J Thromb Haemost. 2016; 14: 1618–28.

28. Doudna JA, Charpentier E. Genome editing. The new frontier of genome engineering with CRISPR-Cas9. Science. 2014; 346: 1258096.

29. Noyes HA, Agaba M, Anderson S, Archibald AL, Brass A, Gibson J, Hall L, Hulme H, Oh SJ, Kemp S. Genotype and expression analysis of two inbred mouse strains and two derived congenic strains suggest that most gene expression is trans regulated and sensitive to genetic background. BMC Genomics. 2010; 11: 361.

30. Speakman JR. Measuring energy metabolism in the mouse - theoretical, practical, and analytical considerations. Front Physiol. 2013; 4: 34.

31. Westbrook R, Bonkowski MS, Strader AD, Bartke A. Alterations in oxygen consumption, respiratory quotient, and heat production in long-lived GHRKO and Ames dwarf mice, and short-lived bGH transgenic mice. J Gerontol A Biol Sci Med Sci. 2009; 64: 443–51.

32. Holzenberger M, Dupont J, Ducos B, Leneuve P, Geloen A, Even PC, Cervera P, Le Bouc Y. IGF-1 receptor regulates lifespan and resistance to oxidative stress in mice. Nature. 2003; 421: 182–7.

33. Even PC, Nadkarni NA. Indirect calorimetry in laboratory mice and rats: principles, practical considerations, interpretation and perspectives. Am J Physiol Regul Integr Comp Physiol. 2012; 303: R459–76.

34. Arch JR, Hislop D, Wang SJ, Speakman JR. Some mathematical and technical issues in the measurement and interpretation of open-circuit indirect calorimetry in small animals. Int J Obes (Lond). 2006; 30: 1322–31.

35. Butler AA, Kozak LP. A recurring problem with the analysis of energy expenditure in genetic models expressing lean and obese phenotypes. Diabetes. 2010; 59: 323–9.

36. Kaiyala KJ, Schwartz MW. Toward a more complete (and less controversial) understanding of energy expenditure and its role in obesity pathogenesis. Diabetes. 2011; 60: 17–23.

37. Kaiyala KJ, Morton GJ, Leroux BG, Ogimoto K, Wisse B, Schwartz MW. Identification of body fat mass as a major determinant of metabolic rate in mice. Diabetes. 2010; 59: 1657–66.

38. Tschop MH, Speakman JR, Arch JR, Auwerx J, Bruning JC, Chan L, Eckel RH, Farese RV, Jr., Galgani JE, Hambly C, Herman MA, Horvath TL, Kahn BB, et al. A guide to analysis of mouse energy metabolism. Nat Methods. 2011; 9: 57–63.

39. Mina AI, LeClair RA, LeClair KB, Cohen DE, Lantier L, Banks AS. CalR: A Web-Based Analysis Tool for Indirect Calorimetry Experiments. Cell Metab. 2018; 28: 656–66 e1.

40. Miller RA, Harper JM, Galecki A, Burke DT. Big mice die young: early life body weight predicts longevity in genetically heterogeneous mice. Aging Cell. 2002; 1: 22–9.

41. Greer KA, Canterberry SC, Murphy KE. Statistical analysis regarding the effects of height and weight on life span of the domestic dog. Res Vet Sci. 2007; 82: 208–14.

42. He Q, Morris BJ, Grove JS, Petrovitch H, Ross W, Masaki KH, Rodriguez B, Chen R, Donlon TA, Willcox DC, Willcox BJ. Shorter men live longer: association of height with longevity and FOXO3 genotype in American men of Japanese ancestry. PLoS One. 2014; 9: e94385.

43. Heiman ML, Tinsley FC, Mattison JA, Hauck S, Bartke A. Body composition of prolactin-, growth hormone, and thyrotropin-deficient Ames dwarf mice. Endocrine. 2003; 20: 149–54.

44. Lusk G. (1917). The elements of the science of nutrition: WB Saunders Company).

45. Leone S, Shohreh R, Manippa F, Recinella L, Ferrante C, Orlando G, Salvatori R, Vacca M, Brunetti L. Behavioural phenotyping of male growth hormone-releasing hormone (GHRH) knockout mice. Growth Horm IGF Res. 2014; 24: 192–7.

46. Facchini FS, Hua N, Abbasi F, Reaven GM. Insulin resistance as a predictor of age-related diseases. J Clin Endocrinol Metab. 2001; 86: 3574–8.

47. Paolisso G, Gambardella A, Ammendola S, D’Amore A, Balbi V, Varricchio M, D’Onofrio F. Glucose tolerance and insulin action in healthy centenarians. Am J Physiol. 1996; 270: E890–4.

48. Dominici FP, Hauck S, Argentino DP, Bartke A, Turyn D. Increased insulin sensitivity and upregulation of insulin receptor, insulin receptor substrate (IRS)-1 and IRS-2 in liver of Ames dwarf mice. J Endocrinol. 2002; 173: 81–94.

49. Dominici FP, Arostegui Diaz G, Bartke A, Kopchick JJ, Turyn D. Compensatory alterations of insulin signal transduction in liver of growth hormone receptor knockout mice. J Endocrinol. 2000; 166: 579–90.

50. Turner AN, Andersen RS, Bookout IE, Brashear LN, Davis JC, Gahan DM, Davis JC, Gotham JP, Hijaz BA, Kaushik AS, McGill JB, Miller VL, Moseley ZP, et al. Analysis of novel domain-specific mutations in the zebrafish ndr2/cyclops gene generated using CRISPR-Cas9 RNPs. J Genet. 2018; 97: 1315–25.

